# Soil Determines Microbial Functionality and Genotype Guides Endophytic Recruitment to Adaptability in Sugarcane Systems

**DOI:** 10.1101/2025.08.30.672968

**Authors:** J.D. Ferreti, Bianca Ribeiro, João de A. Bonetti, Luis Eduardo A. Camargo, S. Creste, E.E. Kuramae, C.B. Monteiro-Vitorello

## Abstract

Soil properties critically shape sugarcane growth and its microbiome, yet their influence on gene expression remains unclear. We investigated the combined effects of soil type (clayey and sandy loam) and sugarcane genotype (IACSP-5503 and IACSP-6007) on microbiome composition and plant transcriptional profiles. Bacterial communities from soils and stalk tissues, as well as transcriptomes of 48-hour sprouted buds grown for 10 months, were analyzed. Results showed that IACSP-5503 (adapted to low-fertility soils) and IACSP-6007 (less adapted) recruited endophytic microbiota in a soil-genotype-dependent manner. In sandy loam, IACSP-5503 promoted diverse plant growth-promoting bacteria (PGPB) (including *Burkholderia*, *Leifsonia*, and *Mycobacterium*), associated with nitrogen fixation, hormone production, and stress tolerance, while IACSP-6007 displayed reduced PGPB diversity and transcriptomic signatures of nutrient deficiencies. Conversely, in clayey soil, IACSP-6007 recruited more PGPBs (such as *Pseudomonas*, *Bacillus* and *Klebsiella*) linked to nutrient acquisition and defense responses. Both genotypes exhibited enhanced expression of defense- and antioxidant-related genes in clayey soil, suggesting priming effects. Overall, our findings reveal soil-dependent, genotype-specific microbial recruitment strategies, including a potential “cry for help” mechanism in IACSP-5503, reflecting adaptation under nutrient-poor conditions. The combined 16S metataxonomic and transcriptome data offered insights into how soil and genotype shape microbial recruitment and transcriptional plasticity in sugarcane.

## 1. Introduction

Historically cultivated for sugar production, sugarcane has gained prominence in recent decades as a leading crop for biofuel production. As its economic significance continues to rise, so does the demand for enhanced productivity achieved through environmentally sustainable practices. However, the widespread cultivation of sugarcane presents sustainability challenges, primarily due to its heightened requirements for water, fertilizers, and pesticides (Vandenberghe et al., 2022). Understanding how sugarcane interacts with several biotic and abiotic factors to improve productivity and essential agronomic practices for developing sustainable agricultural production in different environments.

Like other plants, sugarcane engages with a diverse microbial community that can be tailored to meet its specific needs (Camargo et al., 2023; Costa et al., 2023). Studies highlight the crucial role of microbial communities associated with various organisms, including plants, animals, and humans, in shaping their health and survival (Banerjee; van der Heijden, 2023). These communities are often referred to as a “second genome” because they provide additional genetic information that complements the host’s genome. Similarly, the term “extended genotype” emphasizes how the genetic influence of these microbes extends beyond the host’s DNA, affecting traits like disease resistance, metabolism, and stress tolerance (Banerjee; van der Heijden, 2023). In terms of agriculture and plant health, soils are the main source of microbiomes which represent the largest microorganism’s biomass fraction on the planet, with emphasis on bacteria that account for almost 15% of the total living biomass, followed by the fungi community representing 2% (Bar-On et al., 2018; Banerjee; van der Heijden, 2022). These soil microbiome communities are essential nutrient cycling, organic matter decomposition, soil structure formation, and carbon storage, influencing the ecosystem health and playing a vital role in carbon regulation and agricultural productivity (Degani et al., 2019; Garland et al., 2021; Ercole et al., 2025). The soil microorganisms are crucial in the biogeochemical cycling of elements like nitrogen, phosphorus, sulfur, and iron, directly impacting plant health (Bünemann et al., 2018; Garland et al., 2021). Furthermore, when addressing soil health, other aspects such as soil tilth, fertility, and quality reflect the physical, chemical, and biological properties that govern soil function (Bünemann et al., 2018).

While indicators such as nutrient cycling and microbial composition are important for assessing soil health, their significance depends on the specific environmental context (Fierer et al., 2021). Plants selectively recruit microorganisms from the rhizosphere that play crucial roles in nutrient acquisition, hormone modulation, and defense against biotic and abiotic stresses (van der Heijden et al., 2008; Bardgett & van der Putten, 2014). In sugarcane, beneficial bacterial genera such as *Burkholderia*, *Pantoea*, *Klebsiella*, *Azospirillum*, *Bacillus*, and *Pseudomonas* have been frequently identified, many of which possess traits associated with nitrogen fixation, phosphate solubilization, production of growth hormones, and synthesis of antimicrobial compounds (Quecine et al., 2012; de Souza et al., 2016; Singh et al., 2021). Additionally, certain rhizosphere microorganisms produce secondary metabolites like ferulic, coumaric, and syringic acids that contribute to plant defense mechanisms (Setiawati & Mutmainnah, 2016). These functional traits underline the potential of the sugarcane-associated microbiome to support plant growth and resilience, particularly under stressful environmental conditions. For example, such microbial interactions have been shown to significantly enhance sugarcane resistance to smut disease, highlighting the role of endophytes in inducing systemic resistance (Mehdi et al., 2025). These insights into microbial interactions within sugarcane open new avenues for research, shedding light on the potential of beneficial microbes to enhance plant growth and resilience.

Therefore, beyond the presence of beneficial microorganisms in the soil, it is essential to understand how plants, particularly at the genotype level, can recruit a microbiome that promotes their growth and development selectively. Genetic subtle variations in plants can significantly influence microbial colonization and its impact on plant health (Haney et al., 2015; Spooren et al., 2024). Research on Arabidopsis and poplar trees has demonstrated that even subtle differences in genotypes, such as mutations in receptor kinase genes or those involved in lignin biosynthesis, can alter the composition of root- and endosphere-associated microbial communities (Haney et al., 2015; Beckers et al., 2016; Song et al., 2021). The genetic diversity of the host plant plays a crucial role in shaping the microbial community, as different varieties and cultivars can release distinct profiles of exudates (Trivedi et al., 2020; Spooren et al., 2024). Also, the exudatés specific composition varies between different plant root zones, plant developmental stages and environmental conditions (López et al., 2023). This specificity in exudate composition acts as a selection mechanism, promoting the growth of specific microorganisms that may benefit from the compounds released depending on the plant genotype and tissue (Pascale et al., 2020; Trivedi et al., 2020).

This study aims to deepen our understanding of the influence of soil-genotype interactions in microbial recruitment, plant health, and growth in sugarcane. Specifically, we examine the effects of two contrasting soil types: sandy loam and clayey, each with distinct physic-hydric and fertility profiles. We focus microbial recruitment and community dynamics in two sugarcane genotypes, particularly their growth and resilience. The selected varieties, IACSP01-5503 and IACSP04-6007, were chosen for their contrasting adaptability: IACSP01-5503 is highly resilient and competitive in restrictive environments with low water retention and limited fertility, while IACSP04-6007 demonstrates strong maturation throughout the harvest season in soils with better water retention. To investigate the specific microbial communities these genotypes recruit in different soil types and plant compartments, we employed 16S rRNA gene amplicon sequencing. We also analyzed the transcriptional profiles of both genotypes’ vegetative organs to assess the influence of soil and microbial communities on gene expression. Through this study, we aimed to provide insights that could guide agricultural practices and inform targeted research to breed optimized sugarcane varieties.

## 2. Methodology

### 2.1 Experimental Design

The experiment was conducted with two varieties grown in two distinct fields characterized by different soil types. The first area, located in Itirapina, São Paulo, Brazil, features sandy loam soil, while the second area, situated in Piracicaba, São Paulo, Brazil, consists of clayey soil. The experimental units were arranged in a completely randomized block design with six replications. The seedlings were planted in December 2022 at both cultivation areas, and the plants were maintained in the field for ten months (**Figure 1**). Each experimental plot consisted of four rows of 4 meters with a planting spacing of 0.5 meters, totaling 32 plants per plot and a total of 192 plants of each variety for each of the two cultivation areas. The chosen varieties were IACSP01-5503, a robust and highly competitive cultivar in restrictive environments, particularly those with low water retention capacity and poor natural fertility (https://www.iac.sp.gov.br/noticiasdetalhes.php?tag=1279), and IACSP04-6007, which demonstrates strong maturation throughout the harvest season and performs better in soils with higher water retention compared to those with low water retention (https://www.novacana.com/noticias/novas-variedades-cana-iac-liberadas-setor-terca-feira-231121). All seedlings provided by Dr. Silvana Creste from the Sugarcane Research Center (IAC/Apta – Ribeirão Preto) were disease free plants produced by *in vitro* culture. For this study, we selected two types of soil: sandy loam soil and clayey soil. The two growing areas were monitored over a 10-month period for precipitation.

**Figure 1.**
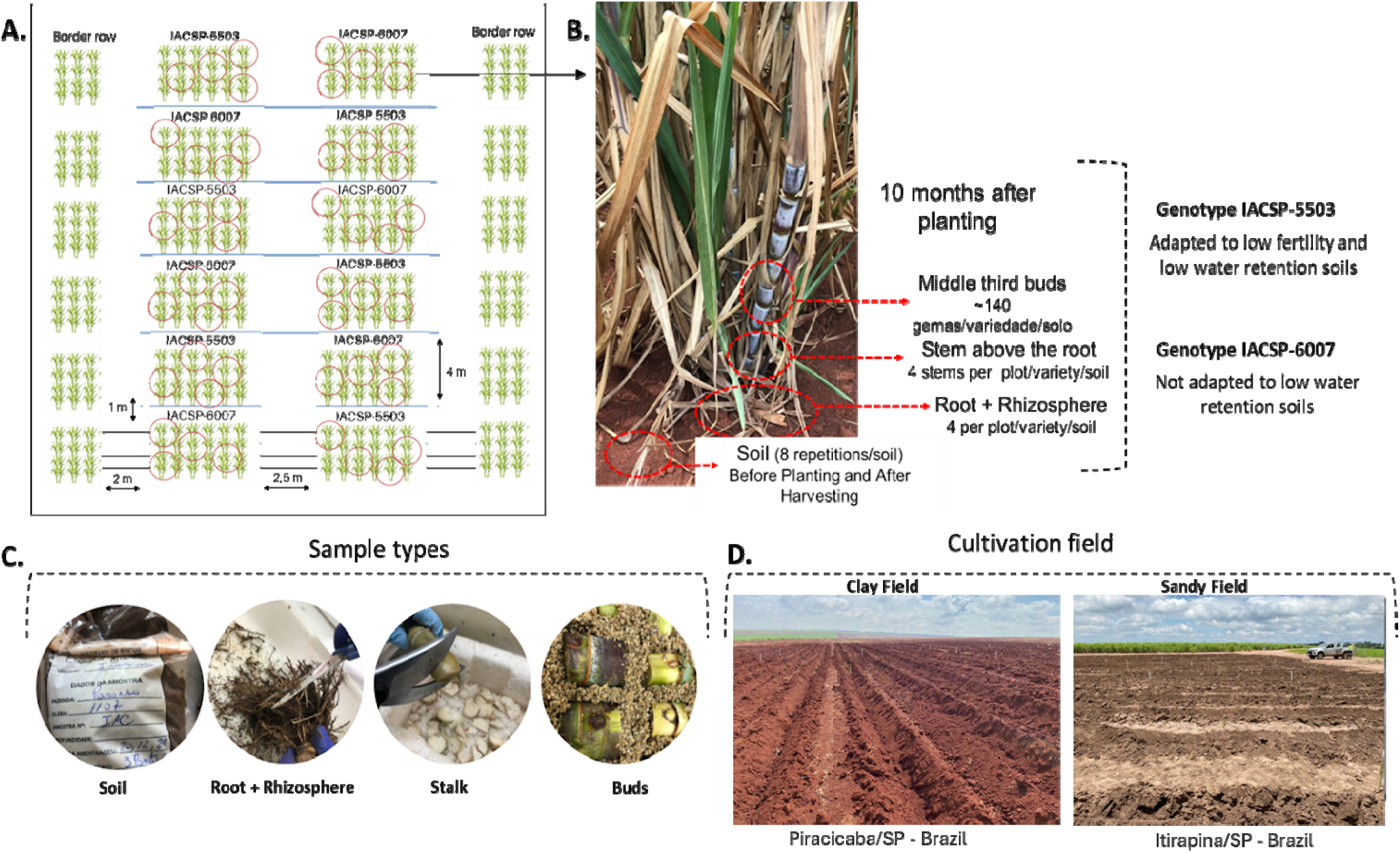
Experimental design and sampling preparation. The experiment used two sugarcane varieties, IACSP01-5503 and IACSP04-6007, grown in sandy and clay soils. **A.** Seedlings were planted in December 2022 at Piracicaba and Itirapina, São Paulo, Brazil, in a randomized block design with six replicates. Each plot consisted of 32 plants, arranged in four rows with 0.5-meter spacing. **B.** Samples were collected ten months after planting (October 2023), with plant material taken from the rhizosphere+roots, stem above the roots, and buds. Soil samples were collected before planting and after harvesting. **C.** Images depict sampling and preparation processes, including washing and bud preparation. **D.** GPS coordinates show the locations of the experimental fields.

### 2.2 Soil physical and chemical analysis

Samples from each of the two experimental areas were collected before planting in December of 2022 for physical and chemical soil analyses and on the day of plant collection in November of 2023 for chemical soil analyses alone. Eight soil samples were collected at both times and composed into four samples. Soil analyses were conducted at the Laboratory of Soil, Plant and Fertilizers Analysis of UNESP/FCA (Faculdade de Ciências Agrárias, Botucatu, SP) to evaluate physical properties, macronutrients (P, K, Ca, S e Mg), micronutrients (B, Cu, Fe, Mn e Zn), pH, and soil organic matter (SOM) (Raij et al., 2001).

### 2.3 Biological material sampling for bacterial 16s rRNA sequencing

Plant material was collected 10 months after planting (MAP) (Figure 1). We collected one randomly selected plant in each row/variety/experiment, thus totaling 24 plants (four plants per plot x 6 plots). Samples of the rhizosphere+root, the stalk just above the root and buds were collected from each plant. In addition, the soil samples were collected at two distinct time points: before planting and after harvest, to assess the initial microbiome conditions and evaluate the changes in the microbiome throughout the growing cycle by metataxonomic analysis. The soil samples were collected from six different points within the experimental fields for microbiome analysis and, like the other samples (root/rhizosphere and stalk), were immediately cooled with liquid nitrogen and stored at −80°C for preservation. Maceration was performed using a CryoMill (Retsch Milling & Sieving), and the macerated samples were kept at −80°C until DNA extraction. The soil samples and the root/rhizosphere and stalk samples were used for DNA extraction and included in the metataxonomy analysis to evaluate the microbial composition and diversity in each sample type collected.

### 2.4 16S rRNA sequencing and bioinformatics analysis

DNA extraction was performed using the PowerSoil DNA Isolation Kit (Mo Bio Laboratories, Inc) for the soil samples and CTAB protocol for plant tissues (Doyle and Doyle, 1990). The DNA was subsequently quantified using a Qubit Fluorometer (Thermo Fisher Scientific). The 16S region of bacterial DNA was sequenced at the Functional Genomics Center, ESALQ, University of São Paulo (Piracicaba/BR), using the Illumina HiSeq 2500 platform. The paired-end sequencing of the V4 region of the 16S gene wasperformed (515FB: GTGYCAGCMGCCGCGGTAA; 806RB: GGACTACNVGGGTWTCTAAT) with 2 x 390 bp reads, according to Parada et al. (2016) and Apprill et al. (2015). PNA clamp sequences were introduced to minimize the amplification of the 16S region of mitochondria and chloroplasts (de Souza et al., 2016).

The raw 16S rRNA sequences were processed using cutadapt v5.1 (Martin, 2011) to remove adapter and primer sequences. Quality filtering was performed with the filterAndTrim function from the DADA2 package v1.37.0 (Callahan et al., 2016) to discard low-quality reads. Error rates were learned, and the dada algorithm was applied to correct sequencing errors. Forward and reverse reads were merged to create a sequence table, and chimeric sequences were removed, and the Amplicon Sequence Variants (ASVs) were used for clustering. The resulting ASV table was imported into a *phyloseq* object (McMurdie & Holme, 2013). The taxonomic assignment was conducted using the Silva reference database (SILVA Release 138.2) (Quast et al., 2013) with the *assignTaxonomy* function in dada2. A rarefaction curve was generated using the vegan package (Oksanem et al., 2015) to evaluate sequencing depth. All bioinformatic analyses were conducted on a high-performance computing cluster.

For the dissimilarity analysis, the dataset was transformed using a centered log- ratio approach to capture the compositional structure of the data (Gloor et al., 2017). Data ordination was performed using Euclidean distance, and principal coordinates were plotted along two axes. Significance was assessed through permutational multivariate analysis of variance (PERMANOVA) with a 5% significance level, utilizing the “adonis” function from the *vegan* package v2.6-10 (Oksanem et al., 2015). Additionally, we identified the most significant genera for group classification using a random forest analysis and estimated the functional abundance with the FAPROTAX algorithm (Sansupa et al., 2021). The Microeco package v1.15.0 (Liu et al., 2021) was used to perform all the final analyses.

### 2.5 Biological material sampling for transcriptome analysis

Buds from the middle third of the stalks of the IACSP-5503 and IACSP-6007 varieties, cultivated in clay and sandy soils, were collected 10 months after planting. Initially, the buds underwent cleaning in running water, followed by thermal treatment (30 minutes at 52 °C) and chemical treatment (10 minutes in a 0.4% m/v solution of sodium hypochlorite) with three successive washes in distilled water. The buds were then placed in vermiculite and kept in the dark at 28°C for 48 hours. The buds were collected in liquid nitrogen for RNA sequencing and stored at −80°C until maceration. The maceration process was conducted using a CryoMill (Retsch Milling & Sieving). RNA extraction followed the Trizol + PureLink Mini Kit (Thermo Fisher Scientific) protocol, and RNA was treated with DNAse I (Sigma Aldrich Corporation). RNA quantification was performed using a NanoDrop 2000 spectrophotometer (Thermo Fisher Scientific), and visualization was done on a 1.5% (w/v) agarose gel. Each RNAseq treatment consisted of three biological replicates, with each replicate composed of 15 buds.

### 2.6 Transcriptome sequence and bioinformatics analysis

The samples were sent to the Functional Genomics Center, ESALQ, University of São Paulo (Piracicaba/BR) for quality control and further processing for library construction using the TruSeq Stranded Illumina kit. Sequencing was performed on a NextSeq (Illumina) instrument with a 2x100bp configuration, with an average coverage of 10 million clusters, meaning 20 million paired reads per sample.

The quality of the paired-end reads was assessed using FastQC v0.10.1. Low-quality bases (Phred < 20), adapter sequences, poly-A tails, and reads shorter than 50 bp were removed using Trimmomatic v0.36 (Bolger et al., 2014). Ribosomal RNA sequences were filtered by mapping the reads against the SILVA NR ribosomal RNA database (SILVA Release 138.2) using the Hisat2 alignment software. For the alignment of the reads obtained after these cleaning steps, the reference complete tetraploid genome of R570 was used (Healey et al., 2024). All alignment steps were performed using Hisat2 v2.1.0. The FeatureCounts software from the Subread package (Liao; Smyth; Shi, 2014) was used to process the BAM mapping outputs from Hisat2 and generate the read count tables. The EdgeR v3.21 software from the Bioconductor package (Robinson; Mccarthy; Smyth, 2010) was used to identify the Differentially Expressed Genes (DEGs). Only genes with CPM (counts per million) values greater than one in all three biological replicates were considered expressed. DEGs were considered statistically significant when the FDR (false discovery rate) was < 0.05 and were represented as Log2 Fold Change (inoculated/control) values. Only genes with Log2 fold change > 1 were retained for further analysis.

### 2.7 Annotation and gene ontology analysis

The BLAST2GO OmicsBox v3.4 was used with the default parameters to assign GO (Gene Ontology) terms to the DEGs. GO enrichment analysis was conducted with the TOPGO in R software using the two-sided Fisher’s exact test with the p-value set at ≤ 0.05. The genes annotated with the same name were represented as one, and the log2 fold change was calculated as the meaning of those values.

## 3. Results

### 3.1 Analysis of physical and chemical soil compositions

Particle-size analysis confirmed distinct natures of the two soils based on textural classification: one classified as sandy loam, containing 833 g kg[¹ of sand, and the other as clayey, with 625 g kg[¹ of clay (**Supplementary File 1 - Table 1**). In the sandy loam soil, the sand fraction consisted of 65% coarse sand (2.0–0.2 mm) and 35% fine sand (0.2–0.05 mm). Chemical analyses of nutrient composition before planting and after harvest for both soil types are presented in **Supplementary File 1 - Table 1**. In general, the clay soil showed medium soil fertility, while the sandy soil exhibited high soil fertility, likely due to the intensive fertilization applied. According to rainfall records, the sandy soil region experienced a two-month period with a more pronounced water deficit (**Supplementary File 1 - Table 1**).

### 3.2 16S rRNA amplicon sequence

#### 3.2.1 DNA sequencing and bioinformatics analysis

Overall, 23,881–85,526 clean 16S rRNA gene sequences per sample were obtained. For alpha-diversity analysis, all samples were rarefied to 23,000 16S rRNA gene sequences and for the other downstream analysis, the sequences were used without rarefaction. The sequencing depth was adequate since the Good’s coverage sequences in all samples were above 99%.

#### 3.2.2 The bacterial community in the two sugarcane production areas before planting and after harvesting

We compared bacterial diversity and taxonomic composition to characterize the soil microbiome before planting and after harvesting. Analysis of 16S rRNA ASVs showed significantly higher alpha diversity in sandy loam than in clayey soils (Dunn’s Kruskal-Wallis test). While no significant differences were found between time points in clayey soil, sandy soil showed a significant shift over time. Overall, ASV-based diversity was consistently greater in sandy soils **(Figure 2A**).

**Figure 2.**
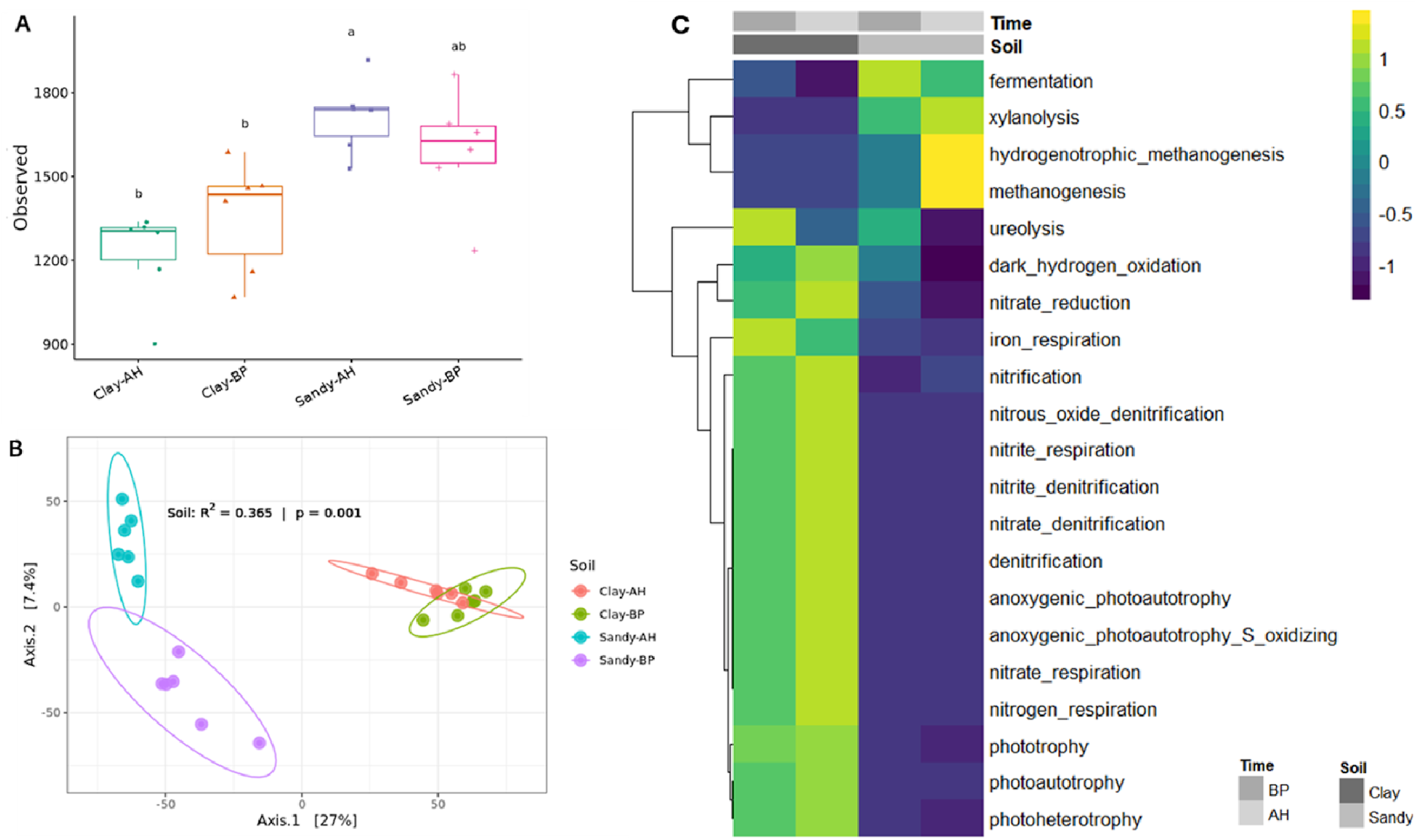
Diversity Analysis of Samples. **A.** Alpha Diversity. Observed diversity, represented by the total number of ASVs, of microbial communities in each soil type (Clay and Sandy) and collection time point (Before Planting -BP and After Harvesting – AH). Significant differences were found between soil types at both time-points. Within the same soil type, significant differences were observed between collection times for sandy soil samples, but no significant differences were detected for clay soil samples (p < 0.05 in Kruskal-Wallis post hoc Dunn test). **B.** Beta Diversity. Overall comparisons of samples based on principal coordinate analysis (PCoA) measured by Euclidean distance. Points closer together represent similar microbial communities, while points farther apart represent dissimilar microbial communities. The statistical significance of differences among both substrates and tree species was calculated by perMANOVA and is shown in the figure. Time-points: Before Planting (BP) and After Harvesting (AH). **C.** Heatmap of Differential Microbial Functions between Clayey and Sandy Soils. Microbial functions were predicted using the FAPROTAX algorithm, and differential abundances between the two soil types were visualized in the heatmap. Statistical analysis was performed using the Kruskal-Wallis test to evaluate overall significance of functional differences between the groups. Post hoc comparisons were conducted using Duncan’s Multiple Range Test to determine which specific soil types exhibited significant differences in microbial functions. Functions were considered differentially abundant if the p-value was less than 0.05.

To assess bacterial community similarity across soil types and time points, principal coordinate analysis (PCoA) based on Euclidean distance was performed (**Figure 2B**). Both soil type and collection time significantly influenced sample separation (PERMANOVA: R² = 0.365, p = 0.001), with soil type showing a stronger effect. The two time points of sandy loam soil showed more distance between themselves than the samples collected from clayey soil. The variance of both soil type patterns was particularly significant, with the first principal component explaining 24% of all differences.

Additionally, at the ASV level, sandy soil consistently showed higher total and unique taxonomic richness than clayey soil at both time points. In sandy loam soil, 5,216 ASVs were detected before planting and 5,193 after harvesting, with 2,627 and 2,949 unique ASVs at each respective time, and 1,091 shared between both. In clayey soil, 3,908 ASVs were found before planting and 3,800 after, with 1,617 and 1,468 unique ASVs, respectively, and 967 shared across time points (**Supplementary File 2- Figure 2A-B**).

The dominant bacterial phyla were similar in both soils, with Actinobacteriota, Proteobacteria, Acidobacteriota, and Firmicutes being the most prevalent before planting and after harvesting. Relative abundances varied: Actinobacteriota (32.2– 34.5% in clayey; 23–21% in sandy), Proteobacteria (21.3–19.9% in clayey; 27.8–22.1% in sandy loam), Acidobacteriota (∼12% in both), and Firmicutes (6.1–8.3% in clayey; 12.7–14% in sandy). At the genus level, *Acidothermus*, *Conexibacter*, and *Gaiella* were more abundant in clayey soil, while *Sphingomonas*, *Bacillus*, and *Solirubrobacter* dominated in sandy loam soil, indicating distinct microbial profiles at this taxonomic level (**Supplementary File 2 - Figure 2)**.

Functional predictions using FAPROTAX identified 54 processes, with 38 differing significantly between soil types and time points (**Supplementary File 2 - Table 2**). Functions related to the nitrogen cycle and carbon/energy pathways (e.g., photoheterotrophy, photoautotrophy, anoxygenic photoautotrophy) were overrepresented in clayey soil. In contrast, methanogenesis, particularly hydrogenotrophic, was more prominent in sandy loam soil (**Figure 2C**), indicating distinct metabolic potential shaped by soil composition.

#### 3.2.3 Contrasting Soil Properties Shape Microbial Potential Functions and Soil Health Dynamics in Sugarcane Development

To evaluate similarities in bacterial communities across sample types (soil, rhizosphere+root, and stalks) and treatments (sugarcane genotypes and soil types), a principal coordinate analysis (PCoA) based on Euclidean distance was conducted. The analysis showed clear distinctions among sample types and between soil types (**Supplementary File 2- Figure 4**). Bacterial communities clustered primarily by soil type, indicating it as the main driver of microbial assemblage across all sample types (PERMANOVA: R² = 0.071, p = 0.001).

The microbiomes of the root+rhizosphere differed from the stalks regardless of plant genotypes and soil types (PERMANOVA: R² = 0.26, p = 0.001 for clay; R² = 0.27, p = 0.001 for sandy soil) (**Figure 3A, 3B**). However, when comparing the root+rhizosphere communities between genotypes within the same soil, no significant differences were observed (clay: R² = 0.1071, p = 0.126; sandy: R² = 0.0805, p = 0.755) (**Figure 3C, 3D**). In contrast, endophytic communities in stalks differed significantly between genotypes in both soils (clay: R² = 0.135, p = 0.002; sandy: R² = 0.2464, p = 0.004), suggesting that bacterial composition in stalks is influenced by both soil type and plant genotype (**Figure 3E, 3F**).

**Figure 3.**
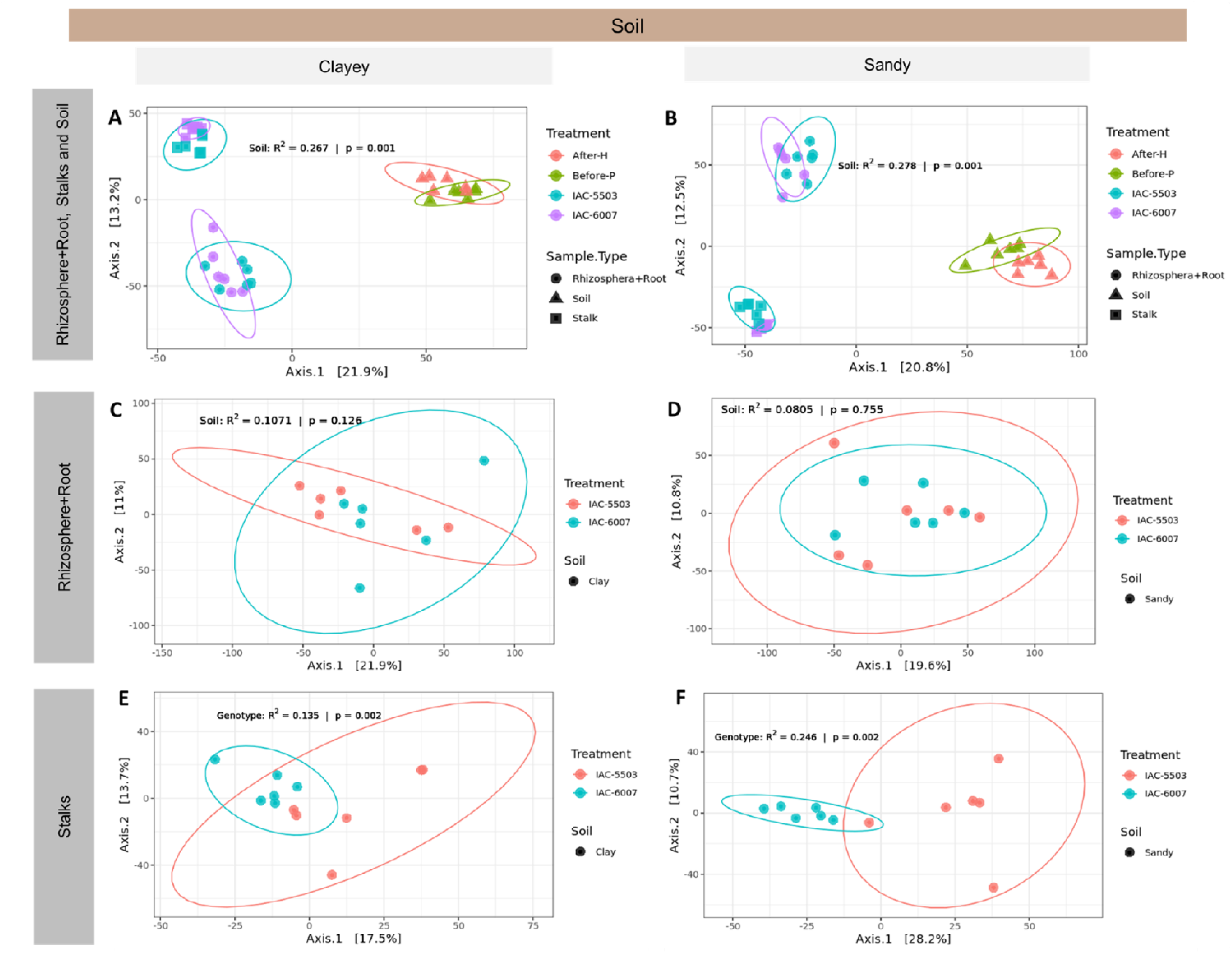
Beta Diversity. Overall comparisons of samples based on principal coordinate analysis (PCoA) measured by Euclidean distance. Points closer to each other represent similar microbial communities, while points farther apart represent dissimilar microbial communities. PCoA analyses of bacterial communities were performed independently for each type of soil. **A** – **B** consider all sample types (soil, rhizosphere + roots, and stalks), while **C** - **D** focus on rhizosphere + root; **E** and **F** solely on stalks for each soil type. **A** and **B** show that sample types are the major driving factors for bacterial community assemblage in both soils, with different shapes indicating the different sample types. **C** and **D** represent the bacterial community in the rhizosphere + root that don’t show significant differences between the genotypes. **E** and **F** show that genotypes drive the bacterial community dissimilarity in stalk samples from both soil origins, with different colors indicating different genotypes.

To investigate differences in stalk-associated microbial communities between genotypes and soil types, a random forest analysis was performed at the genus level (**Figure 4**). Based on a Kruskal-Wallis test, 88 taxa were identified as significantly represented in these compartments (**Supplementary File 2 - Table 1**). In clayey soil, the IACSP-6007 genotype was associated with *Pseudomonas*, *Paenibacillus*, *Klebsiella*, *Pseudarthrobacter*, *Bacillus*, and *Paenarthrobacter*. In contrast, *Kitasatospora* was identified as a unique biomarker for IACSP-5503 in the same soil. Sandy loam soil yielded a greater number of biomarkers for both genotypes. For IACSP-5503, marker genera included *Burkholderia*-*Caballeronia*-*Paraburkholderia*, *Terriglobus*, *Mycobacterium*, *Leifsonia*, *Luteibacter*, *Acidisoma*, *Gryllotalpicola*, and *Acidibacter*. In the same soil, IACSP-6007 was associated with *Acinetobacter*, *Neorhizobium*, *Methylobacterium*-*Methylorubrum*, *Amnibacterium*, and *Klenkia*.

**Figure 4.**
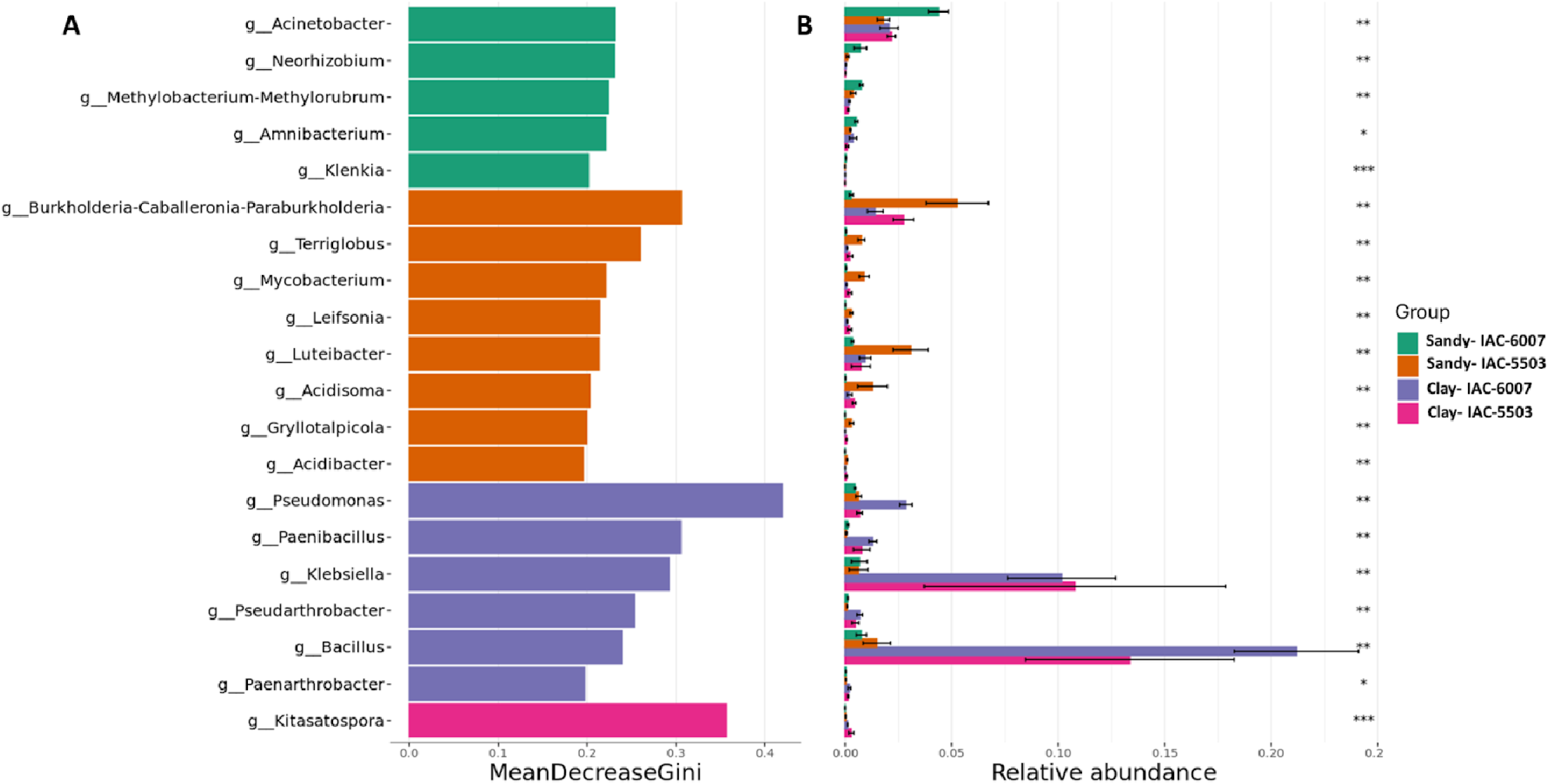
Random Forests Analysis for Biomarkers of Each Genotype (IACSP-5503 and IACSP-6007) in Stalk Samples from Two Different Soils (Sandy – S; Clay - C). **A.** The 20 most important microbial genera for the classification of each group. **B.** The relative abundance of each of these genera in each soil (Sandy and Clay) and genotype (IACSP-5503 and IACSP-6007). Data are presented as mean ± standard deviation.

### 3.3 Genotype-Specific Transcriptional Responses to Soil Type Reveal Differential Plasticity in Sugarcane

#### 3.3.1 Differential gene expression analysis

To investigate differential gene expression (DEGs) in the IACSP-5503 and IACSP-6007 genotypes under different soil conditions, we first assessed genotype performance by comparing gene expression between IACSP-5503 and IACSP-6007 within each soil type (sandy loam and clayey). In clayey soil, 7,806 genes were downregulated, and 5,994 were upregulated. In sandy loam soil, fewer genes were differentially regulated, with 2,458 downregulated and 2,704 upregulated (log2FC>1). In addition, to evaluate how soil type modulates gene expression within each genotype, we compared their expression profiles between sandy loam and clayey soils. For IACSP-5503, 9,622 genes were downregulated and 9,289 were upregulated. In contrast, IACSP-6007 exhibited fewer differentially regulated genes, with 2,718 downregulated and 3,918 upregulated. The DEG results are represented in Venn diagrams, illustrating the number of genes identified as differentially expressed in each comparison (**Figure 5**).

**Figure 5.**
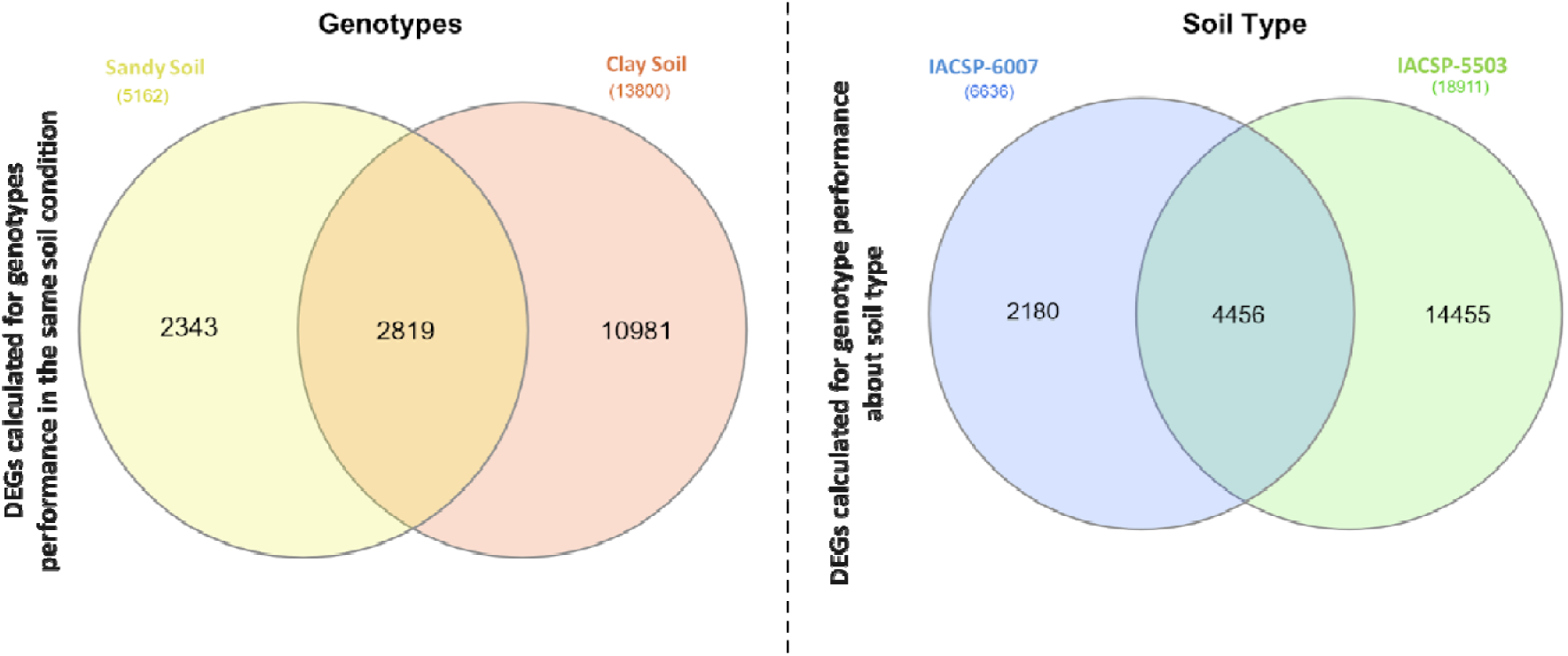
Differential expression gene analysis of IACSP-5503 (IAC5503) and IACSP-6007 (IAC6007) genotypes from sandy loam and clayey soils. Genotypes. - To evaluate genotype performance under the same soil conditions, we calculated the differentially expressed genes (DEGs) by comparing gene expression between the two genotypes (IACSP-5503 and IACSP-6007) within the same soil type. **Soil Type** - To assess how genotype performance varies across different soils, we calculated the DEGs by comparing gene expression of each genotype between the sandy and clay soils. Venn diagrams display the number of genes identified as differentially expressed (DEGs) using EdgeR software from the Bioconductor package (Robinson; McCarthy; Smyth, 2010). The intersections of the identified DEGs from the mapping results were considered for further analysis.

#### 3.3.2 Enrichment Analysis of GO Terms

The molecular processes driving the sugarcane response to different microbiome and soil conditions were investigated by gene ontology (GO) enrichment analysis of DEGs. The enrichment analysis of the IAC-5503 and IAC-6007 varieties DEGs grown in sandy loam and clayey soils showed contrasting molecular responses based on the soil type. In sandy loam soil, cell wall growth mechanisms were up-regulated, while in clayey soil both varieties present several hormone and stress related responses up-regulated, for both genotypes (**Figure 6, Supplementary File 2 – Figures 5-6**).

**Figure 6.**
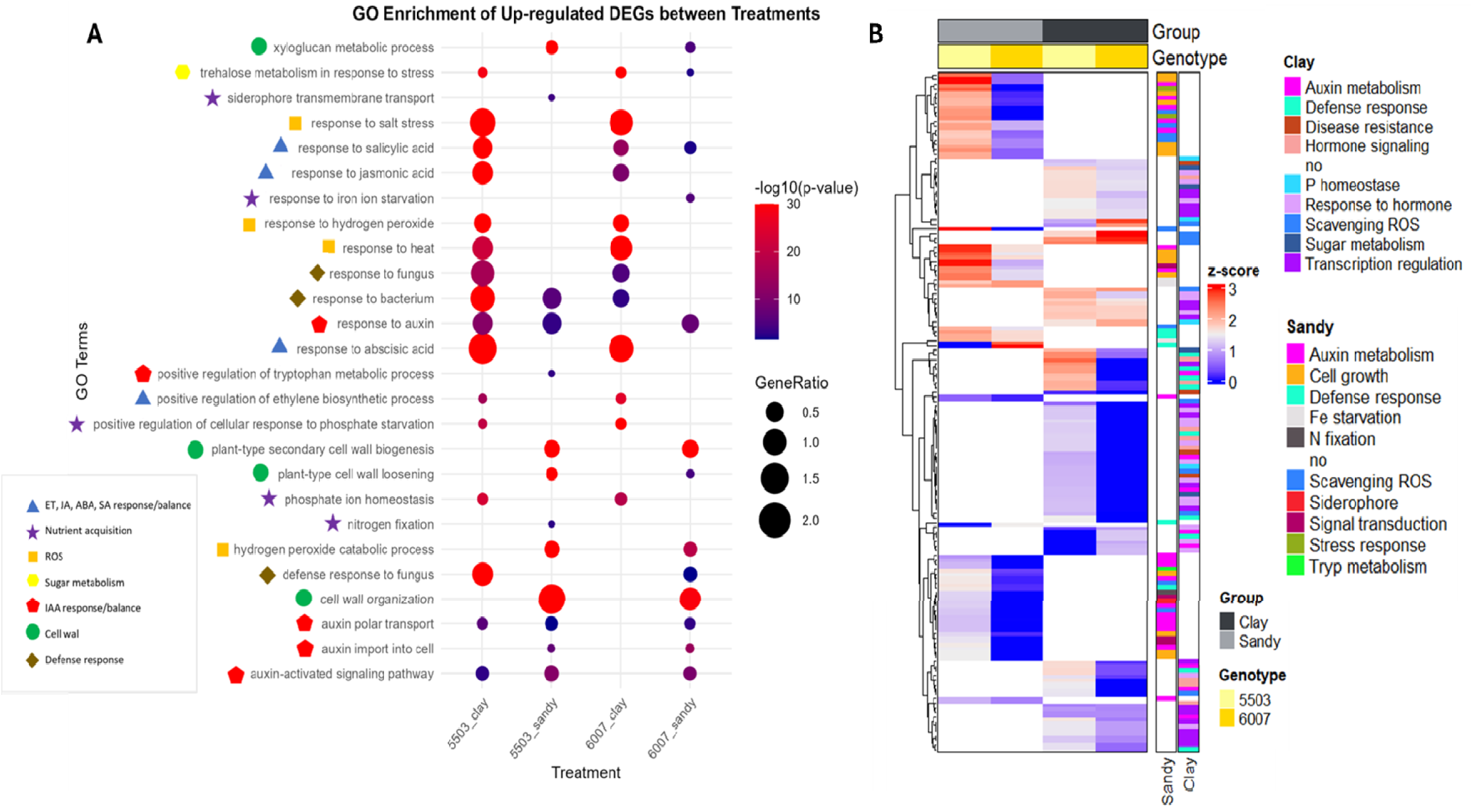
Functional categorization and expression patterns of upregulated differentially expressed genes (DEGs) across sugarcane genotype–soil combinations. **A**. Bubble plot representing Gene Ontology (GO) enrichment analysis of upregulated DEGs based on biological process terms, with a significance threshold of *p* ≤ 0.05. The y-axis lists significantly enriched GO terms, while the x-axis shows the different genotype–soil treatments. Bubble size reflects the proportion of upregulated genes associated with each term (GeneRatio), and color represents statistical significance (–log□□(*p*-value)). GO terms were manually grouped into functional categories and are represented by distinct shapes and colors to facilitate interpretation. **B.** Hierarchical clustering heatmap showing z-score normalized expression profiles of selected upregulated DEGs included in the GO enrichment analysis for each treatment (panel A). Genes are annotated with the functional categories used in the bubble plot, represented by colored bars on the right. Columns represent individual samples organized by genotype and soil type. Both genes and samples were clustered using hierarchical clustering to reveal expression patterns across treatments.

##### 3.3.2.1 Genotypes transcriptional profile in sandy loam soil

In sandy loam soil, Gene Ontology (GO) enrichment analysis revealed specific terms for the IACSP-5503 and IACSP-6007 genotypes (**Figure 6A**). The IACSP-5503 genotype showed enrichment for “nitrogen fixation,” accompanied by the upregulation of genes related to ammonium assimilation, such as glutamine synthetase-2 (**Supplementary File 2 – Figure 5**). In contrast, the IACSP-6007 genotype did not exhibit this pattern.

Regarding iron acquisition, the non-adapted IACSP-6007 genotype displayed enrichment of the GO term “response to iron ion starvation,” while the adapted IACSP-5503 genotype showed enrichment for “siderophore transmembrane transport,” with increased expression of genes important for siderophore uptake. In addition, investigation of indole-3-acetic acid (IAA)-related genes revealed enrichment of GO terms associated with IAA production and transport, including “auxin import into cell,” “auxin polar transport,” “auxin-activated signaling pathway,” and “response to auxin” in both genotypes (**Figure 6A**). Upregulation of auxin-activated enzymes was observed, particularly in the IACSP-5503 genotype. Additionally, the GO term “positive regulation of tryptophan metabolic process” was enriched in both varieties, characterized by the upregulation of the gene encoding the enzyme related to WAT1 (Walls Are Thin1) (**Supplementary File 2 – Figure 5**).

Concerning plant defense mechanisms, the IACSP-6007 genotype grown in sandy loam soil exhibited enrichment of the GO term “defense response to fungus,” while the IACSP-5503 genotype showed enrichment for “response to bacteria”. Both genotypes displayed enrichment of the GO term “hydrogen peroxide catabolic process,” with upregulation of enzymes related to hydrogen peroxide degradation, more pronounced in the adapted genotype (**Figure 6 AB**).

Overall, the heatmap analysis (**Figure 6B**) for sandy loam soil revealed distinct patterns of gene expression across functional categories for both genotypes. For IACSP-5503, categories such as nitrogen fixation, siderophore transmembrane transport, and auxin metabolism showed prominent upregulation. For IACSP-6007, categories like response to iron ion starvation and defense response to fungus were notably enriched. Both genotypes showed significant activity in categories related to cell growth and wall organization, including “plant-type cell wall loosening,” “cell wall organization,” “xyloglucan metabolic process,” and “plant-type secondary cell wall biogenesis” (**Figure 6A**), accompanied by elevated expression of various enzymes associated with plant cell growth, such as celluloses, xyloglucans, and expansins (**Supplementary File 2 – Figure 5**).

##### 3.3.2.2 Genotypes transcriptional profile in clayey soil

In clayey soil, Gene Ontology (GO) enrichment analysis revealed specific terms for both IACSP-5503 and IACSP-6007 genotypes (**Figure 6A**). An enrichment of “positive regulation of cellular response to phosphate starvation” and “phosphate ion homeostasis” was observed in both varieties. This was accompanied by the upregulation of genes known to enhance phosphorus use efficiency, such as purple acid phosphatase (PAP) and genes containing the Syg1/Pho81/XPR1 (SPX) domain (**Supplementary File 2 - Figure 6**). In parallel with nutrient acquisition, GO terms linked to defense responses were also enriched. Both genotypes displayed enrichment of “response to fungus” and “response to bacteria”, with “defense response to fungus” being specifically enriched in IACSP-5503.

The defense signature observed was further complemented by hormonal and redox-related responses. GO terms enriched in both varieties when grown in clayey soil included “response to hydrogen peroxide”, “response to abscisic acid”, “response to salicylic acid”, “response to jasmonic acid” and “positive regulation of ethylene biosynthetic process” (**Figure 6A**). For “response to hydrogen peroxide”, several heat shock proteins were more up-regulated in the IACSP-6007 variety, while other genes encoding ROS scavenging enzymes such as catalase3 were up-regulated in the IACSP-5503 (**Supplementary File 2 - Figure 6**). Most of these enzymes were also present in the enriched GO terms “response to heat” and “response to salt stress” for both varieties. For “response to abscisic acid” GO term, upregulation of transcription factors (TFs) was observed, mostly in the IACSP-5503 variety. Likewise, most of these TFs were also present in other enriched GO terms such as “response to salicylic acid”, “response to jasmonic acid” and “positive regulation of ethylene biosynthetic process”. The “trehalose metabolism in response to stress” GO term was also enriched in both varieties with an up-regulation of trehalose-6-phosphate synthase gene (TPS) (**Supplementary File 2 - Figure 6**).

In summary, the heatmap analysis (**Figure 6B**) for clayey soil revealed distinct patterns in terms of gene expression level across functional categories for both genotypes. In general, for both IACSP-5503 and IACSP-6007, categories such as phosphate homeostasis, defense responses, and various stress responses (e.g., hydrogen peroxide, heat, salt stress, abscisic acid, salicylic acid, jasmonic acid, ethylene biosynthetic process, and trehalose metabolism) showed prominent upregulation. Specific differences in gene expression within these categories were observed between the two genotypes, as detailed above.

## 4. Discussion

Sustainable plant production depends on habitat provision, soil biodiversity, and efficient nutrient cycling, all of which operate as an interdependent system (Lehmann et al., 2020; Garland et al., 2021; Banerjee & van der Heijden, 2023). Within this context, plant diversity and long-term vegetation cover impact soil health, shaping the microbial communities (Bardgett & van der Putten, 2014; Garland et al., 2021). Therefore, a holistic approach considering vegetation cover (Garland et al., 2021), soil management (Norris & Congreves, 2018; de Graaff et al., 2019), soil physicochemical properties (George et al., 2019; Lehmann et al., 2020), and microbial biomass (McDaniel et al., 2014; Vukicevich et al., 2016) is essential to enhance soil functionality and ensure long-term agricultural sustainability.

### 4.1 Contrasting Soil Types Drive Bacterial Potential Functions and Soil Health Dynamics in Sugarcane Systems

To assess the influence of soil type on sugarcane microbial recruitment, we compared sandy loam and clayey soils under long-term cultivation and two sugarcane genotypes. The two soils differed in texture and chemical characteristics, shaped by management practices. Despite sandy soil being naturally less fertile, fertilization in our experiment resulted in higher measured fertility than clay soils (Cantarella et al., 2022; Samuel & Dines, 2023). Still, sandy soil showed lower water/nutrient retention, reduced carbon accumulation, and drought exposure over time (Garbowski et al., 2023). Also, our data reflected the two-month period when pluviometry data showed distinct values, where the sandy soil was exposed to a prolonged drought. Although both soils shared similar dominant bacterial phyla, the sandy soil showed slightly higher relative abundances of microbial groups positively associated with productivity, such as Proteobacteria and Firmicutes (Romero et al., 2024). Sandy loam soils showed higher methanogenesis potential, particularly via hydrogenotrophic pathways. Although generally more aerated, their low water/nutrient retention can promote localized anaerobic microenvironments (Garbowski et al., 2023), within organic matter aggregates, where methanogenic microbes thrive (Angle et al., 2017).

Conversely, the clay soil’s superior moisture retention likely sustained a microbial functional stability under stress (Romero et al., 2024). The clayey soil offers a more stable microhabitat, favoring bacteria in nitrogen and carbon cycling by preserving organic matter, creating a favorable environment for photoheterotrophic and photoautotrophic microorganisms (Pagliai & Vignozzi, 2002; Lehman et al., 2020). We observed that soil physical structure influences microbial composition and function, consistent with its known impact on nutrient cycling and soil health (George et al., 2019; Lehmann et al., 2020). A relevant component of soil health is resilience, the capacity to maintain functional stability over time despite environmental disturbances (Lehmann et al., 2020). In our study, shifts in bacterial community composition over the 10-month crop cycle revealed greater microbial stability in clayey than in sandy soil. Often, microbial resilience is associated with greater organic matter content and the more stable physical structure typical of clayey soils (Bonetti et al., 2017; Bucka et al., 2019).

### 4.2 Endophytic Microbiome Composition in Sugarcane Is Driven by Genotype and Modulated by Soil Type

Soil is the primary microbial reservoir for sugarcane plants (Bulgarelli et al., 2013; de Souza et al., 2016), where root exudates, complex organic compound mixtures varying by species, genotype, and developmental stage, structure soil and rhizosphere communities, influencing endophyte recruitment (Trivedi et al., 2020; Spooren et al., 2024). Our experiment used plants regenerated from micromeristematic tissues, which ensured they were free of associated microorganisms before field cultivation. We found no genotype effect on root plus rhizosphere microbiome within the same soil, but apparent varietal differences in stalk endophytic communities. The absence of a roots genotype effect reflects strong soil and plant developmental stage influence at sampling, consistent with Chaparro et al. (2014), who showed rhizosphere communities converge at similar growth stages.

Differences in soil type reflect distinct environmental conditions that may influence both the selection and persistence of plant growth-promoting bacteria (PGPB) associated with sugarcane. These bacteria contribute to enhanced plant stress tolerance by potentially modulating physiological and biochemical processes, including phytohormone production (e.g., indole-3-acetic acid, IAA), nutrient acquisition (such as phosphate solubilization and siderophore production), ethylene regulation through ACC deaminase activity, and the activation of antioxidant defense systems (Ercole et al., 2025). Consistent with these microbial functions, our RNA-seq data revealed soil-type-dependent transcriptional reprogramming in sugarcane, with enrichment of gene ontology terms linked to these same physiological processes, varying according to the genotype and the specific soil conditions.

#### Sandy loam soil

##### Microbiome composition and nutrient acquisition

IACSP-5503 harbored a stalk-associated community enriched in putative PGPB in sandy loam soil. Consistent with IACSP-5503 outperforming IACSP-6007 in sandy loam soils, its microbiome was enriched for *Burkholderia, Luteibacter*, *Mycobacterium*, and *Leifsonia*, genera with reported roles in nitrogen fixation, phytohormone production, and adaptation to low nutrient environments (Naveed et al., 2014; Xu et al., 2023). In *Miscanthus floridulus*, *Leifsonia* and *Burkholderia* participate in ammonium/nitrate transformations, underscoring nitrogen cycling functions (Zhaoyu et al., 2024). IACSP-5503 also showed higher *Acidibacter* abundance, which, though not a nitrogen fixer, alters soil pH, influencing nitrogen availability (Falagán & Johnson, 2014). In line with these features, only IACSP-5503 showed enrichment of the GO term nitrogen fixation in the host transcriptome, suggesting coupling between microbial functions and host gene expression in sandy loam soil.

##### Iron homeostasis and siderophores

IACSP-6007 showed enriched “response to iron ion starvation,” consistent with reduced iron bioavailability, possibly due to lower *Acidibacter* abundance. Acidibacter-mediated pH modulation, via ferric iron reduction, can shift iron speciation and bioavailability (Falagán & Johnson, 2014). By contrast, IACSP-5503’s stalk endophyte community was enriched in siderophore-producing genera (*Burkholderia*, *Leifsonia*, *Mycobacterium*, *Luteibacter*), enhancing iron acquisition. Although *Acinetobacter* and *Methylobacterium*, also siderophore producers, were more abundant in IACSP-6007, the transcriptome indicated more effective uptake in IACSP-5503, including enriched siderophore transmembrane transport GO term and higher ABC transporter B family gene expression.

##### Hormonal signaling and growth

RNA-seq analysis revealed that both sugarcane genotypes displayed enriched terms related to auxin biosynthesis and transport, but auxin-activated gene expression was markedly higher in IACSP-5503 under sandy loam conditions. The stronger hormonal response was accompanied by the upregulation of WALLS ARE THIN1 (WAT1), a gene involved in auxin/tryptophan metabolism (Ranocha et al., 2013). In contrast, IACSP-6007 showed lower expression of auxin-activated enzymes, suggesting weaker auxin signaling. These transcriptional differences are consistent with the microbial communities associated with each genotype. *Burkholderia*, more abundant in IACSP-5503, as well as *Mycobacterium*, *Luteibacter*, and *Leifsonia*, are well-known producers of both ACC deaminase and IAA. By reducing ACC levels, ACC deaminase lowers ethylene biosynthesis and its inhibitory effects on cell proliferation, while microbial IAA supports root development and stress tolerance (Spaepen & Vanderleyden, 2011; Chieb & Gachomo, 2023). In contrast, *Acinetobacter* and *Methylobacterium*, predominant in IACSP-6007, also produce ACC deaminase and IAA, but their presence did not translate into a comparable transcriptional response.

##### Defense and induced systemic resistance

Both sugarcane genotypes upregulated numerous defense-related genes in sandy loam soil, but the response was consistently stronger in IACSP-5503. This genotype showed enhanced expression of cellulose synthases, chlorophyllase, PR proteins (β-1,3-glucanases, peroxidases, thaumatins), ricin B-like lectins, thaumatin-like proteins (TLPs), as well as ERECTA and AAA ATPases. These proteins contribute to structural reinforcement, oxidative stress mitigation, and resistance to both abiotic and biotic stresses. The upregulation of ricin B-like lectins, associated with drought and osmotic stress tolerance (Sahid et al., 2020), and TLPs, with antifungal and abiotic stress functions (Liu et al., 2010), underscores the convergence of stress-response pathways in IACSP-5503. ERECTA, which encodes a receptor-like kinase implicated in cell wall integrity and defense, further highlights the broader defense potential of this genotype, since mutants lacking this gene are known to be more susceptible to bacterial and fungal pathogens (Shpak et al., 2013). In contrast, IACSP-6007 activated a more limited defense response, with β-1,3-glucanase being the most notable defense-related transcript. These differences suggest that while PGPB, such as *Burkholderia*, can trigger induced systemic resistance (ISR) through the accumulation of PR proteins and secondary metabolites (Singh et al., 2021), the host genotype largely determines the extent and breadth of transcriptional defense activation.

##### Cell wall remodeling and growth

Both sugarcane genotypes activated a transcriptional program associated with cell wall remodeling and growth regulation in sandy loam soil. Enriched processes included cell wall loosening, organization, xyloglucan metabolism, and secondary cell wall biogenesis. Consistently, transcripts for cellulose synthases, expansins, and enzymes of the xyloglucan pathway were upregulated in both genotypes. This conserved response, which sustains cell expansion and mitigates water loss under sandy soil conditions, reflects a canonical stress adaptation mechanism previously described in plants (Han et al., 2015; Marowa et al., 2016; Yang et al., 2020).

#### Clayey soil

##### Microbiome composition and nutrient acquisition

The two genotypes in clayey soil showed apparent genotype-dependent differences in stalk-associated microbiomes. IACSP-6007 stalks were enriched for PGPB genera like *Pseudomonas*, *Bacillus*, *Pseudarthrobacter*, *Paenibacillus*, *Paenarthrobacter*, and *Klebsiella*, which are functionally linked to IAA synthesis, phosphate solubilization, ammonia/siderophore production, and nitrogen fixation (Ercole et al., 2025). Notably, *Pseudomonas*, *Bacillus*, and *Paenibacillus* in IACSP-6007 are recognized for their ability to trigger ISR, potentially enhancing host defense in clayey soils (Ercole et al., 2025). In contrast, IACSP-5503 showed enrichment mainly for *Kitasatospora*, which also possesses plant growth-promoting traits but with a narrower functional profile.

Sugarcane genotypes showed activation of phosphate starvation response pathways, with IACSP-5503 distinguished by the upregulation of SPX4, a central regulator of phosphate signaling. Both genotypes showed enrichment of the GO terms *“positive regulation of cellular response to phosphate starvation”* and *“intracellular phosphate ion homeostasis”*, accompanied by increased expression of PAP genes and the regulators SPX1 and SPX3. These enzymes enhance phosphorus use efficiency by mineralizing organophosphorus compounds (Bhadouria & Giri, 2022; Xu et al., 2022). In Arabidopsis, for instance, AtSPX1 induces PAP2 expression under both phosphate-sufficient and -deficient conditions, whereas AtSPX3 and AtSPX4 act as modulators of feedback regulation (Duan et al., 2008). The SPX homologs in apple are strongly induced under phosphate limitation but repressed when PGPRs are present, showing the microbial contribution to phosphate acquisition (Kural et al., 2024). In this context, we may suggest that phosphate-solubilizing PGPB may further support host nutrition by converting insoluble phosphate into bioavailable forms through extracellular enzymes, organic acid secretion, and siderophore production (Da Silva et al., 2023; Ughamba et al., 2025).

Also, both sugarcane genotypes showed enrichment of the GO term *“trehalose metabolism in response to stress”*, with upregulation of TPS, the enzyme that converts trehalose-6-phosphate (T6P) into free trehalose. Elevated sucrose stimulates T6P synthesis, while increased T6P feeds back to regulate sucrose production and utilization, promoting biomass accumulation and may indicate stress, activating stress-responsive pathways (Eh et al., 2024; Paul et al., 2018). Microbial contributions may reinforce this metabolic adjustment. Certain PGPB, such as *Klebsiella* sp., enhance wheat growth under abiotic stress by expressing trehalose biosynthesis genes (Zhang et al., 2017). Likewise, exogenous application of a T6P precursor increased T6P levels, sucrose accumulation, and endogenous T6P synthesis in wheat (Griffiths et al., 2016). Together, these findings suggest that clayey soil, regardless of genotype, favors T6P metabolism, supporting hormone biosynthesis, stress tolerance, and growth.

##### Hormonal signaling and growth

In clayey soil, both sugarcane genotypes upregulated transcription factors, including WRKY, ERF/AP2, MYB, bZIP, and NAC, associated with hormone-mediated defense priming, but the response was stronger in IACSP-5503. These TFs are frequently reported in plants influenced by PGPB (Kaleh et al., 2024).

##### Defense and induced systemic resistance

Clayey soil fostered beneficial plant-microbe interactions, priming the host against biotic stress. Both genotypes upregulated defense-related genes, including R genes, PAMP-induced protein A70 (PAMP A70), and PR genes (chitinases, PR 1, TLPs). Plant immune perception involves PRR- mediated PAMP-triggered immunity (PTI) (RLKs, RLPs) and NLR R proteins mediating effector-triggered immunity (ETI) (Sekhwal et al., 2015). Increased expression of receptors and R-like genes in clayey soil suggests activated recognition-based immune pathways and a primed immune state. PR 1 was upregulated in IACSP-5503, rapidly induced upon pathogen perception and associated with increased resistance (Han et al., 2023). Chitinases were upregulated in both genotypes, degrading fungal cell wall chitin and induced by salinity, drought, and cold, contributing to broad stress tolerance (Dana et al., 2006; Vaghela et al., 2022).

##### Reactive oxygen species and antioxidant systems

Several PGPB possess catalase activity, detoxifying ROS (Gómez-Godínez et al., 2022). Both genotypes in clayey soil converged on a similar antioxidant transcriptional program, with upregulated catalases, heat shock proteins, and glutathione metabolism enzymes. By contrast, in sandy loam soil, this antioxidant signature was stronger in IACSP-5503 than IACSP-6007, aligning with IACSP-5503’s superior performance. This suggests genotype-specific fine-tuning of antioxidant responses based on soil conditions, with IACSP-5503 showing a more robust response under challenging sandy loam conditions, contributing to enhanced resilience and productivity.

The overall transcriptional landscape in clayey soil indicates a coordinated molecular response to nutrient availability and stress, mediated by plant-microbiome interaction. These findings underscore the importance of considering host genotype and soil type in understanding plant adaptation and developing strategies for sustainable agriculture.

## 5. Conclusion

The results presented in this study provide valuable insights into the complex interplay between plant genotype, soil type, and the associated microbial communities in sugarcane. In summary, our results demonstrate that sugarcane genotype exerts a primary influence over the composition and functional potential of endophytic communities, particularly in internal plant compartments such as stalks. Soil type modulates this effect by shaping the microbial pool available for recruitment and influencing plant physiological responses. The differential recruitment of plant growth-promoting bacteria, along with transcriptomic changes in hormonal signaling, nutrient acquisition, stress tolerance, and redox balance, suggest that genotype-specific microbiota contribute to plant adaptability under contrasting soil conditions (**Figure 7**).

**Figure 7.**
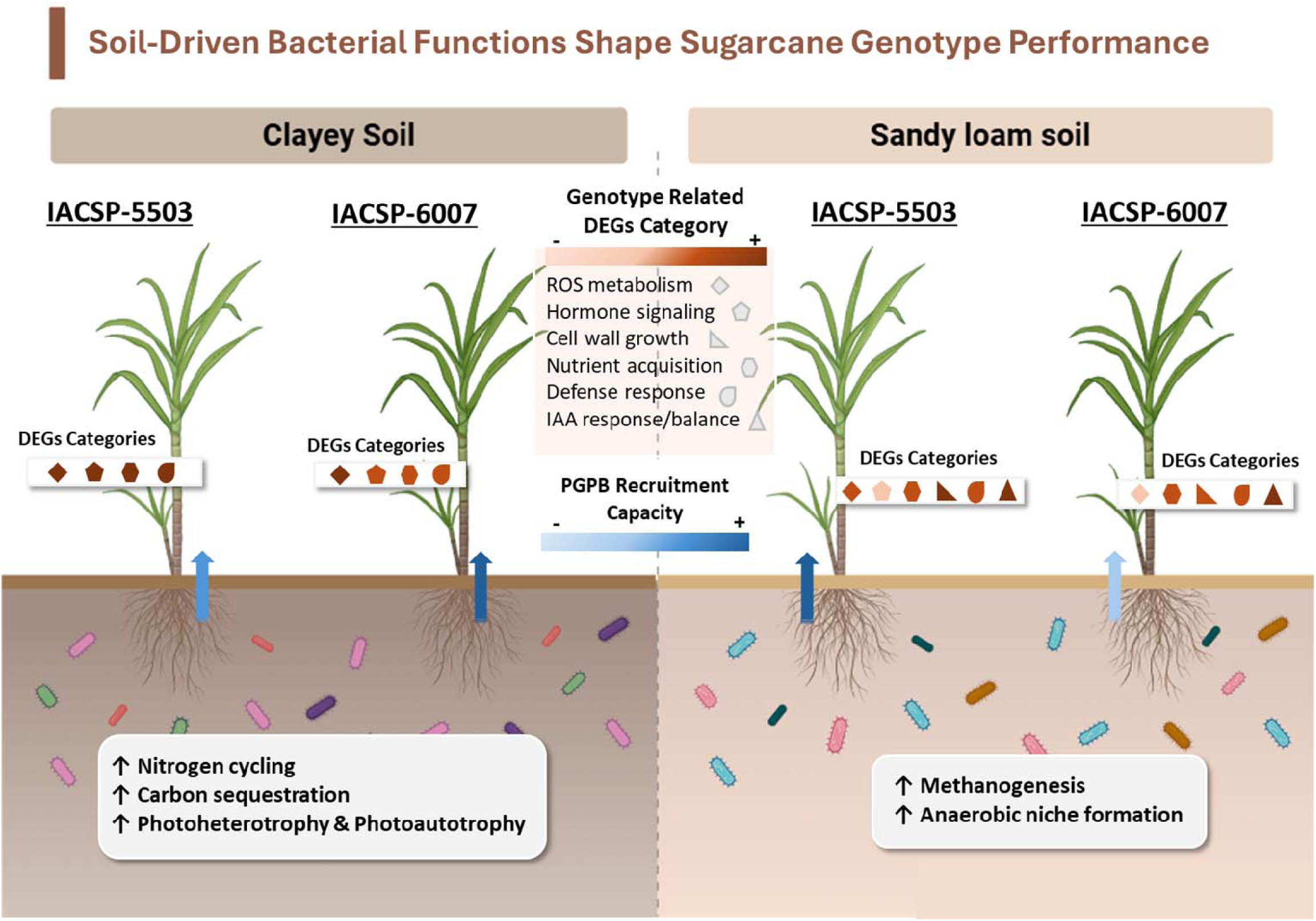
Graphical abstract illustrating how soil type and sugarcane genotype shape bacterial recruitment, microbial functions, and host transcriptional responses. Colored arrows and gradient-filled boxes represent the recruitment capacity of each genotype (IACSP-5503 and IACSP-6007) in clayey and sandy soils. White boxes summarize microbial functions enriched in each soil type. Symbols near each plant represent categories of differentially expressed genes (DEGs), with their shading intensity indicating the magnitude of transcriptional investment. In clayey soils, microbial communities are enriched in carbon and nitrogen cycling functions. Sandy soils harbor stress-tolerant microbes adapted to low-nutrient, low-oxygen conditions. In sandy soil, IACSP-5503 (better adapted) recruits more plant growth-promoting bacteria (PGPB) and activates growth-related genes, suggesting a “cry-for-help” strategy. In clayey soil, IACSP-6007 (less adapted) shows increased PGPB recruitment, potentially compensating for lower genetic potential. The figure highlights genotype-specific microbial recruitment and transcriptional profiles as key drivers of sugarcane adaptation to contrasting soil environments.

These findings emphasize that plant performance under stress is not solely a genotype function but emerges from dynamic interactions with beneficial microorganisms. Understanding genotype-specific microbial recruitment offers opportunities to integrate beneficial microbiome traits into breeding programs, enhancing sugarcane adaptation to specific soils and stress conditions. Multi-omics approaches can support microbiome engineering, through strategies such as synthetic microbial communities and host-mediated selection, to improve stress resilience and overall crop performance. Long-term studies will be crucial to understanding how these interactions evolve over time and across seasons, contributing to more resilient and sustainable sugarcane production systems.

## Declarations

### Ethics approval and consent to participate

Not applicable.

### Adherence to national and international regulations

Not applicable.

### Consent for publication

Not applicable.

### Availability of data and material

The datasets generated and/or analyzed during the current study are available under the NCBI BioProject PRJNA1309163. Additional data can be found in supplementary information files. The complete bioinformatics tools are available in https://github.com/JoyceFerreti/Sugarcane_Microbiome.git and https://github.com/JoyceFerreti/Sugarcane_Microbiome2.git.

### Authors’ contributions

C.B.M.-V. conceived the study and edited the manuscript final version. J.D.F. performed the experiments, analyzed the data, and wrote the manuscript. B.R. analyzed the data and wrote the manuscript. S.C. contributed with plant materials. S.C., J.B., L.E.A.C and E.E.K. provided scientific expertise and manuscript editing. All authors reviewed and approved the final version of the manuscript.

## Supporting information

Soil characteristics and precipitation data

Microbial composition and gene expression

16S amplicon sequencing summary

Transcriptome summary and differential expression

GO terms annotation

## Acknowledgements

Elaine Vidotto Batista for technical support.

## Supplementary Information

**Additional file 1 (.docx):** Soil characteristics and Precipitation data recorded.

**Additional file 2 (.docx):** Microbial Composition Across Samples and Gene expression analysis.

**Additional file 3 (.csv):** 16S amplicon sequencing data summary and taxonomic structure.

**Additional file 4 (.csv):** Transcriptome data summary and differential expression results.

**Additional file 5 (.csv):** GO terms annotation and genes selections.

