## Supplementary material for "Soil Determines Microbial Functionality and Genotype Guides Endophytic Recruitment to Adaptability in Sugarcane Systems": Soil characteristics and precipitation data

**Supplementary Material S1 - Soil characteristics and Precipitation data recorded**

**Table 1. Particle Size Analysis and Soil Textural Classification.**

| Soil Texture Classe | Coarse Sand | Fine Sand | Total Sand | Silt | Clay |
| --- | --- | --- | --- | --- | --- |
|  | g/kg ------------------------------------------------------------------------- | | | | |
| Clay | 52 | 118 | 169 | 206 | 625 |
| Sand Loam | 540 | 293 | 833 | 16 | 152 |

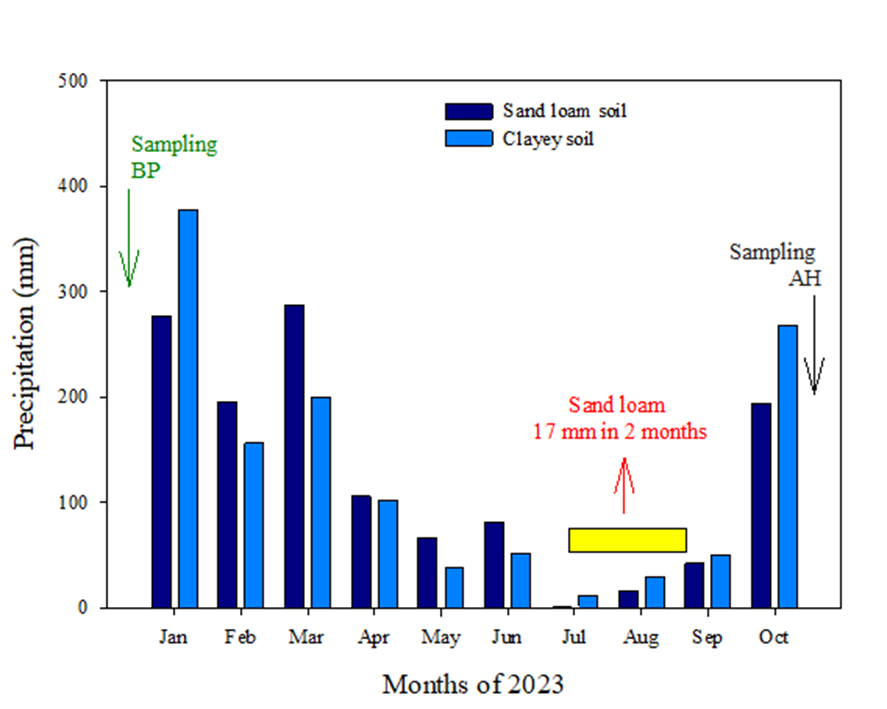

**Supplementary Figure 1.** Cumulative monthly precipitation recorded during the experiment in Corumbataí, SP (sand loam soil) and Iracenápolis, SP (clay soils).

**Supplementary Table 2. Chemical characterization of soil macronutrients and micronutrients - clay soils and sand loam soil.**

| Macronutrients | | | | | | | | | | | | | | | | |
| --- | --- | --- | --- | --- | --- | --- | --- | --- | --- | --- | --- | --- | --- | --- | --- | --- |
| Soil | Time | pH | | SOM | P_resina_ | S | K | Ca | | Mg | Al | | H+Al | SB | CEC | V |
|  |  | CaCl | | g kg^-3^ | mg.dm^-3^-------------- | | mmol_c_.kg^-1^-------------------------------------------------------------------------------------------- | | | | | | | | | % |
| Clay | Planting | 5.10^±0.16^ | | 35.1^±3.55^ | 19.2^±0.68^ | 153.3^±30.7^ | 2.48^±0.41^ | 33.77^±7.17^ | | 19.16^±4.03^ | 0.68^±0.68^ | | 38.20^±3.69^ | 55.4^±11.4^ | 93.6^±8.7^ | 57.5^±6.82^ |
|  | Harvesting | 5.25^±0.36^ | | 24.8^±1.71^ | 32.1^±1.47^ | 78.27^±17.9^ | 1.35^±0.25^ | 28.68^±0.68^ | | 19.34^±5.48^ | 0.14^±0.14^ | | 35.56^±6.07^ | 49.4^±13.65^ | 84.9^±7.94^ | 55.6^±9.39^ |
| Sand Loam | Planting | 5.98^±0.18^ | | 18.0^±1.35^ | 93.4^±19.7^ | 13.0^±3.9^ | 1.12^±0.07^ | 54.91^±7.64^ | | 26.97^±6.27^ | 0.00^±0.00^ | | 14.63^±1.14^ | 83.0^±13.7^ | 97.6^±12.6^ | 83.9^±2.97^ |
|  | Harvesting | 6.69^±0.36^ | | 15.8^±0.80^ | 109.2^±30.5^ | 4.52^±0.26^ | 0.59^±0.03^ | 74.0^±24.9^ | | 45.68^±20.61^ | 0.00^±0.00^ | | 9.83^±0.57^ | 120.3^±45.54^ | 130.1^±45.2^ | 90.2^±2.30^ |
| Micronutrients | | | | | | | | | | | | | | | | |
| Soil | Time | | B | | | Cu | | | Fe | | | Mn | | | Zn | |
|  |  |  | mg/dm^3^------------------------------------------------------------------------------------------------------------------------------------------------------------------------ | | | | | | | | | | | | | |
| Clay | Planting | | 0.55^±0.04^ | | | 3.07^±0.09^ | | | 19.46^±0.74^ | | | 17.62^±1.11^ | | | 1.38^±0.21^ | |
|  | Harvesting | | 0.39^±0.03^ | | | 3.07^±0.16^ | | | 16.01^±1.72^ | | | 14.45^±1.93^ | | | 1.10^±0.08^ | |
| Sand Loam | Planting | | 0.38^±0.11^ | | | 0.90^±0.16^ | | | 59.27^±5.07^ | | | 2.14^±0.57^ | | | 1.65^±0.15^ | |
|  | Harvesting | | 0.22^±0.02^ | | | 0.90^±0.07^ | | | 45.80^±3.85^ | | | 1.28^±0.20^ | | | 1.55^±0.13^ | |

SOM: soil organic matter; BS.: base saturation; SB: sum of bases; CEC: cation exchange capacity; V: percentage of base saturation.
