## Supplementary material for "Soil Determines Microbial Functionality and Genotype Guides Endophytic Recruitment to Adaptability in Sugarcane Systems": Microbial composition and gene expression

**Supplementary Material S2 - Microbial Composition Across Samples and Gene expression analysis**

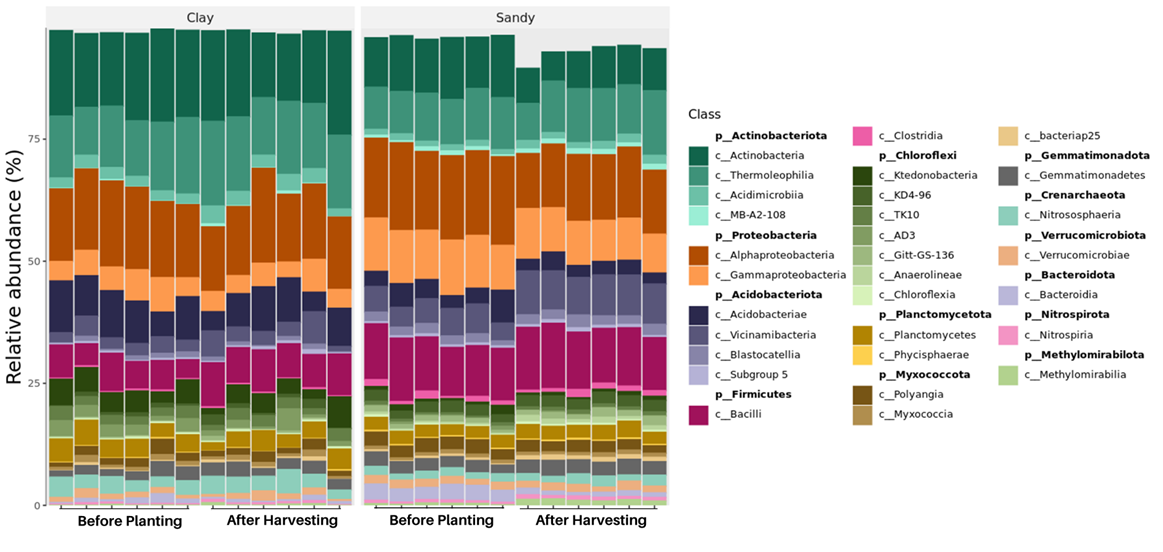

**Figure 1.** Relative abundance of dominant bacterial classes in clay and sandy soils, comparing samples collected before planting and after harvesting. Each bar represents an individual replicate, and classes are color-coded by phylum. Only classes with a relative abundance >1% in at least one sample are shown.

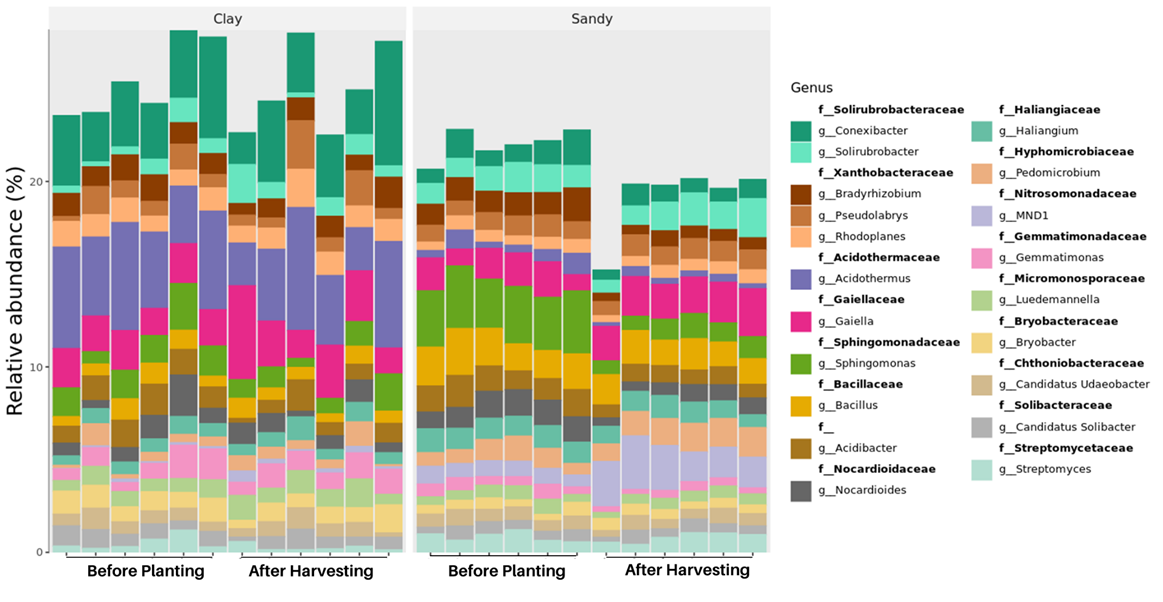

**Figure 2.** Relative abundance of the most abundant bacterial genera in bulk soil samples collected before planting and after harvesting under clay and sandy soil conditions. Each bar represents a replicate, and colors correspond to specific genera grouped by family. Only genera with a relative abundance >1% in at least one sample are displayed.

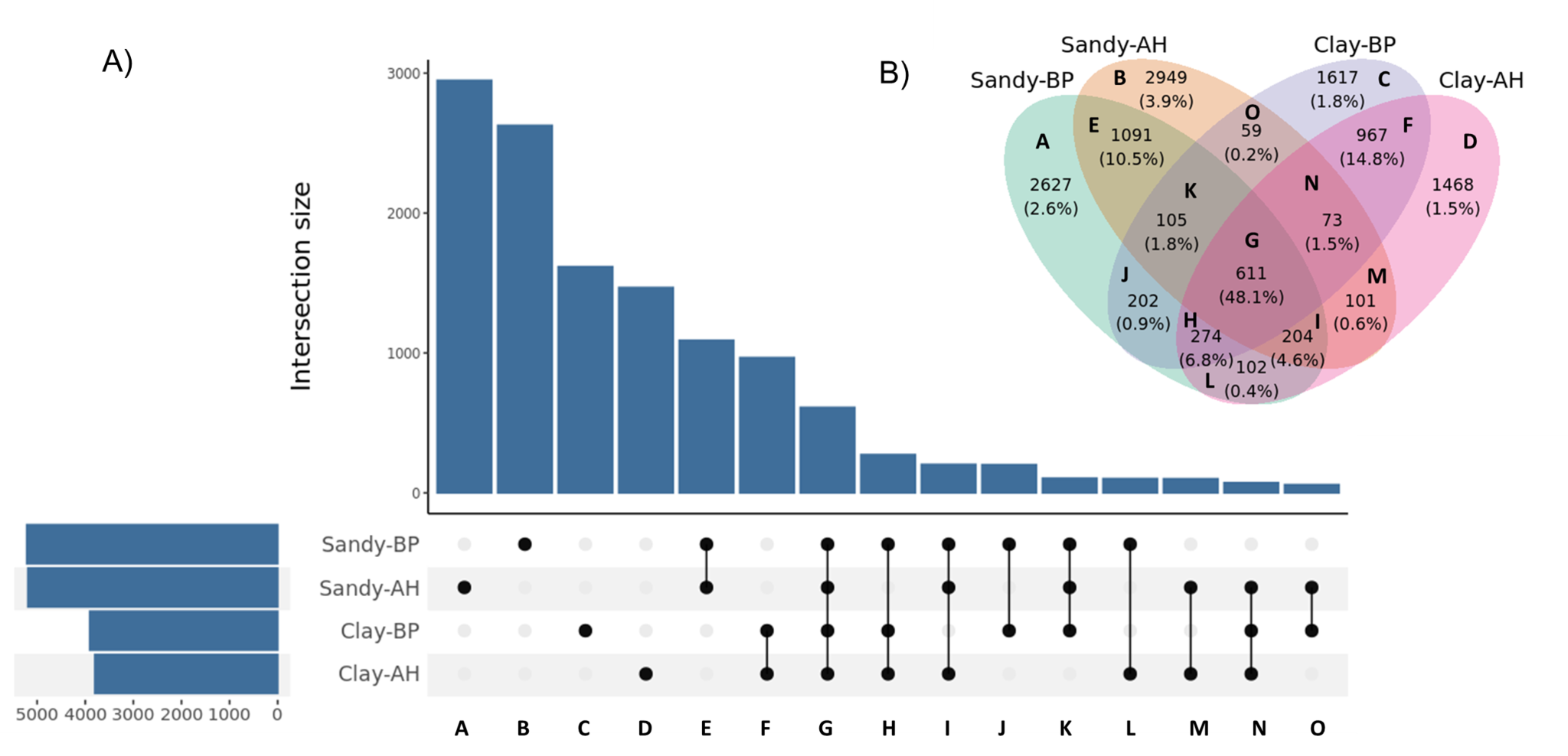

**Figure 3. A.** UpSet plot illustrating the intersection number of the ASVs between the soil types and their collection times. **B.** Venn-Diagram analysis of bacterial community at ASV level for the sandy and clay soils, before planting and after harvesting. The integer is ASV number and the percentage data is the sequence number/total sequence number.

**Table 1.** Microbial functions were predicted by the FAPROTAX algorithm. Differences in microbial function abundances across different soil groups were analyzed using Kruskal-Wallis to assess overall significance. Duncan’s Multiple Range Test was performed as a post hoc analysis to identify which specific groups differed from each other. Differences were considered significant if the p-value was less than 0.05.

| **Function** | **Clay-BP** | **Clay-AH** | **Sandy-BP** | **Sandy-AH** |
| --- | --- | --- | --- | --- |
| anoxygenic_photoautotrophy | a | a | b | b |
| anoxygenic_photoautotrophy_S_oxidizing | a | a | b | b |
| dark_hydrogen_oxidation | a | a | ab | b |
| denitrification | a | a | b | b |
| fermentation | bc | c | a | ab |
| hydrogenotrophic_methanogenesis | b | b | ab | a |
| iron_respiration | a | ab | b | b |
| methanogenesis | b | b | ab | a |
| nitrate_denitrification | a | a | b | b |
| nitrate_reduction | ab | a | bc | c |
| nitrate_respiration | a | a | b | b |
| nitrification | a | a | b | ab |
| nitrite_denitrification | a | a | b | b |
| nitrite_respiration | a | a | b | b |
| nitrogen_respiration | a | a | b | b |
| nitrous_oxide_denitrification | a | a | b | b |
| photoautotrophy | a | a | b | b |
| photoheterotrophy | a | a | b | b |
| phototrophy | a | a | b | b |
| xylanolysis | b | b | a | a |

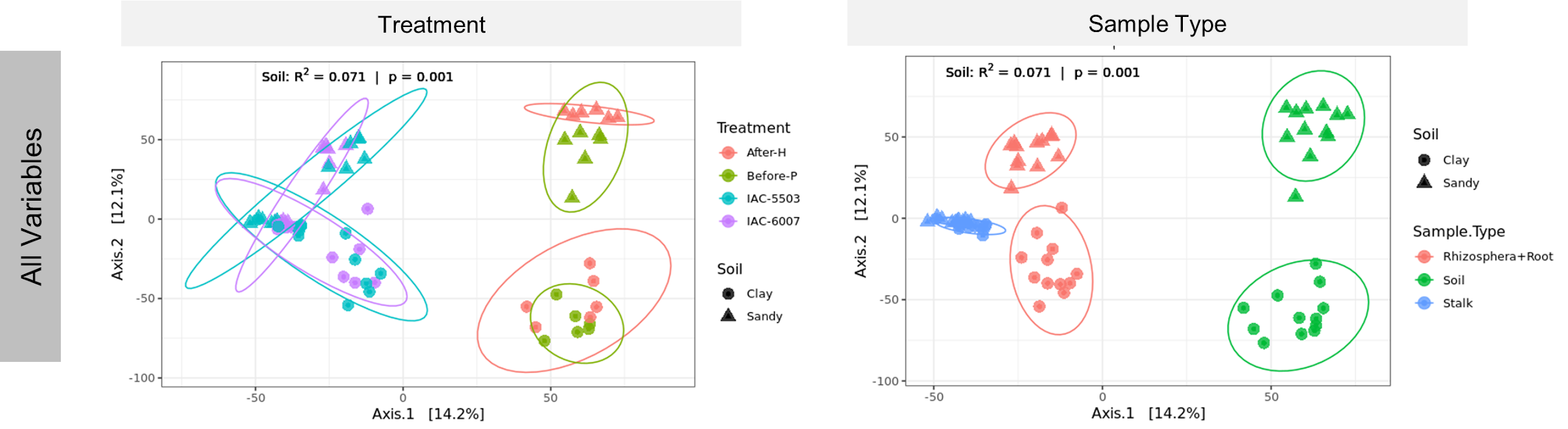
**Figure 4. Beta Diversity.** Overall comparisons of samples based on principal coordinate analysis (PCoA) measured by Euclidean distance. Points closer to each other represent similar microbial communities, while points farther apart represent dissimilar microbial communities. PCoA analyses of bacterial communities were performed independently for groups of samples (soil, rhizosphere + roots, and stalks) with two soil types and two plant genotypes, initially considering "all variables."

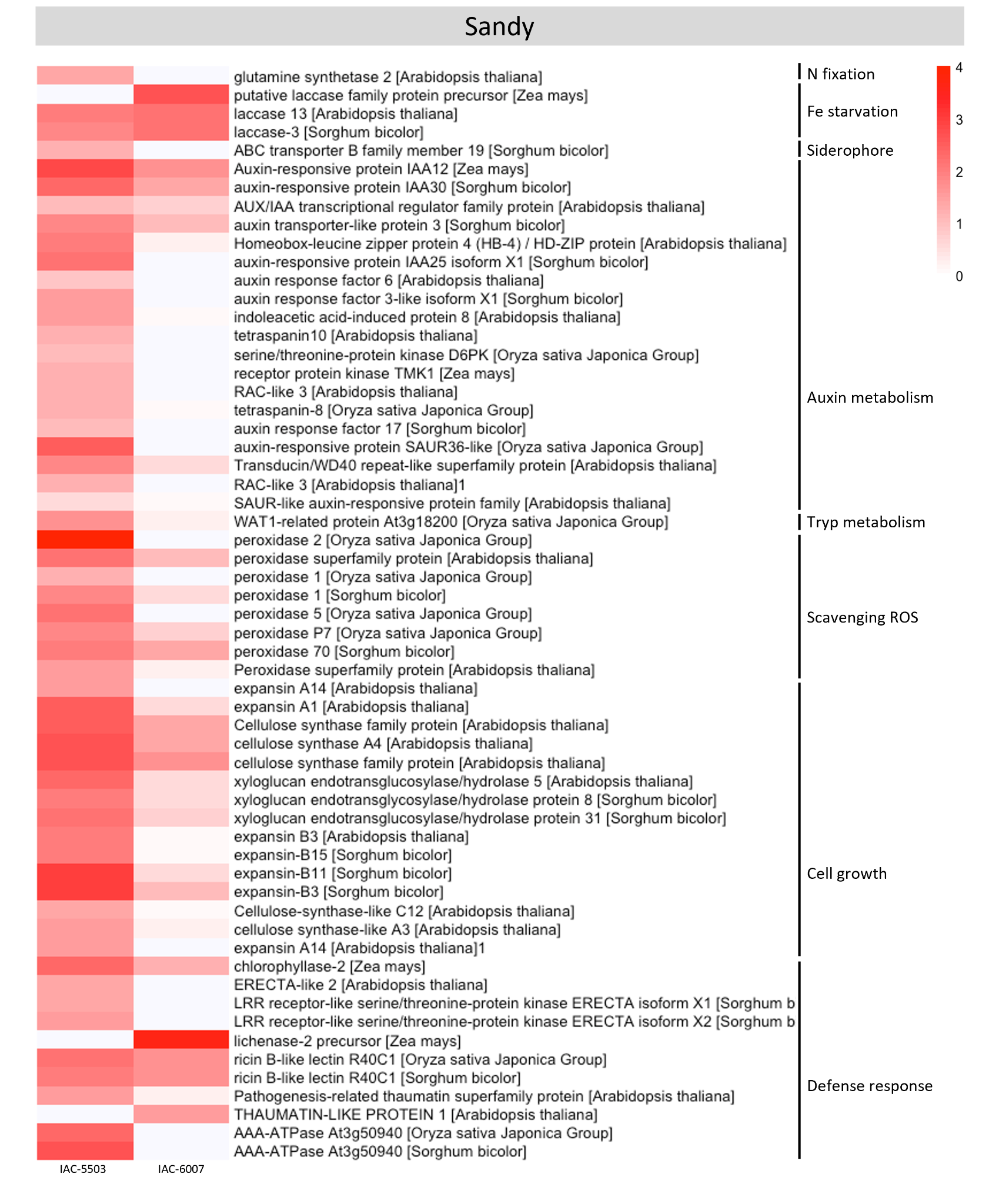

**Figure 5.** Sugarcane DEGs up-regulated in sandy soil. Expression profile of genes related to biological functions represented as values of a log2 fold change (sandy/clay). The heatmap was constructed using the R software package. The red squares represent up-regulated genes which are grouped based on their biological processes.

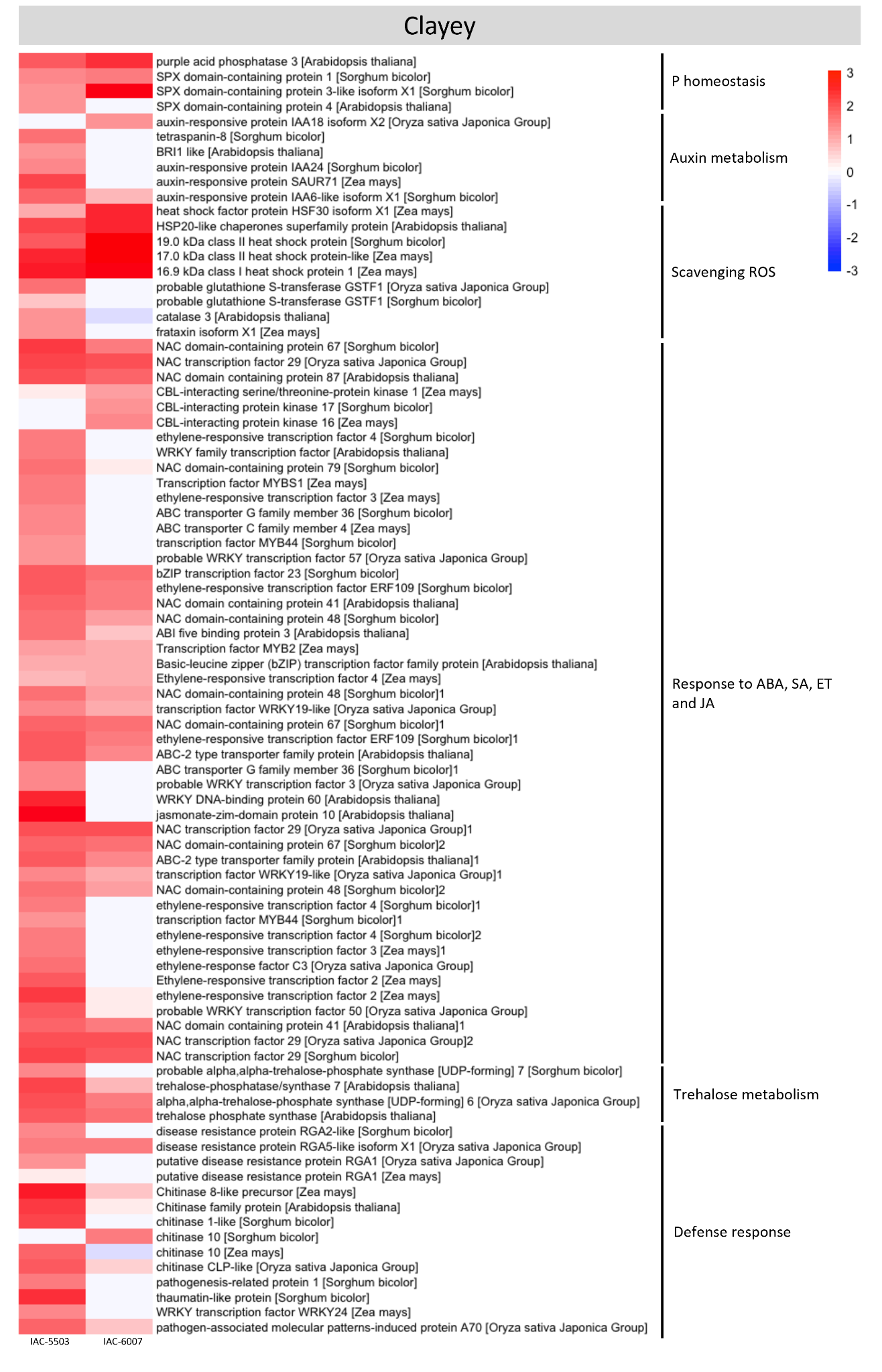

**Figure 6.** Sugarcane DEGs up-regulated in clay soils. Expression profile of genes related to biological functions represented as values of a log2 fold change (clay/sandy). The heatmap was constructed using the R software package. Blue squares represent down-regulated genes, and red squares represent up-regulated ones. The genes are grouped based on their biological processes.
